# Bayesian network models identify co-operative GPCR:G protein interactions that contribute to G protein coupling

**DOI:** 10.1101/2023.10.09.561618

**Authors:** Elizaveta Mukhaleva, Ning Ma, Wijnand J. C. van der Velden, Grigoriy Gogoshin, Sergio Branciamore, Supriyo Bhattacharya, Andrei S. Rodin, Nagarajan Vaidehi

## Abstract

Cooperative interactions in protein-protein interfaces demonstrate the interdependency or the linked network-like behavior of interface interactions and their effect on the coupling of proteins. Cooperative interactions also could cause ripple or allosteric effects at a distance in protein-protein interfaces. Although they are critically important in protein-protein interfaces it is challenging to determine which amino acid pair interactions are cooperative. In this work we have used Bayesian network modeling, an interpretable machine learning method, combined with molecular dynamics trajectories to identify the residue pairs that show high cooperativity and their allosteric effect in the interface of G protein-coupled receptor (GPCR) complexes with G proteins. Our results reveal a strong co-dependency in the formation of interface GPCR:G protein contacts. This observation indicates that cooperativity of GPCR:G protein interactions is necessary for the coupling and selectivity of G proteins and is thus critical for receptor function. We have identified subnetworks containing polar and hydrophobic interactions that are common among multiple GPCRs coupling to different G protein subtypes (Gs, Gi and Gq). These common subnetworks along with G protein-specific subnetworks together confer selectivity to the G protein coupling. This work underscores the potential of data-driven Bayesian network modeling in elucidating the intricate dependencies and selectivity determinants in GPCR:G protein complexes, offering valuable insights into the dynamic nature of these essential cellular signaling components.

## Introduction

### Dynamics play a crucial role in regulating G protein coupling with GPCRs

G-protein-coupled receptors (GPCRs) form a superfamily of membrane proteins of critical importance to cellular function, and the largest group of drug targets (1). Binding of agonists activates the receptor to couple to one or more members of the G protein family in the intracellular region, with different coupling strengths (2). Several three-dimensional structures of GPCR:G protein complexes provide information on the residues that interact at the interface. However, these static structures do not conclusively establish which of these residue interactions are key hotspots for selective coupling of GPCRs with G proteins. Dynamics of the GPCR:G protein complexes play a critical role in regulating selective versus promiscuous coupling (3–7). The structural mechanism(s) of selective or promiscuous G protein coupling by an agonist-GPCR pairing has been the focus of several studies (4–25). Many of the computational studies using MD simulations typically use prior knowledge of the GPCR:G protein systems to decipher the selectivity hotspot residues in the interface. Currently, there is no unbiased data-driven analysis of the dynamics of the interface to decipher the residue hotspots in the interface.

There are three thermodynamic properties of the residue pairs interacting at the GPCR:G protein interface that play an important role in coupling: (i) enthalpy of the interaction (6,7), (ii) spatiotemporal persistence of the interacting residue pairs (5), and (iii) cooperativity resulting from inter-dependencies among the residue pairs (26–30). These three factors are critical to the overall lifetime or the stability of the GPCR:G protein complex and are modulated by the dynamics of the GPCR:G protein interface. In our previous studies, we have shown that both enthalpy and spatiotemporal frequency of the GPCR:G protein interface interactions calculated from the molecular dynamics (MD) simulation trajectories play a crucial role in both selective and promiscuous coupling of the G proteins (5–7). Although cooperativity is known to be important for protein-protein interactions and has been studied in the context of protein folding (28), there is no systematic method to identify such cooperative interactions in GPCR:G protein coupling (29–31). The role of cooperativity among interacting residue pairs is best delineated by protein network properties. Therefore, in this study we have used an interpretable probabilistic Bayesian network (BN) (32,33) model combined with MD simulation trajectories to decipher the role of cooperativity of the GPCR:G protein interactions. Delineating the residue pairs that play a critical role in cooperativity of the GPCR:G protein interactions will lead to identification of hotspot residues in the interface which, in turn, will pave the way to engineer the G protein selectivity preference in GPCRs (4,7). Importantly, we demonstrate the use of BN modeling, an unsupervised data-driven interpretable machine learning method, as a promising approach to identify hotspot residues in protein-protein interfaces.

### Applying Computational Systems Biology / Bayesian networks to mine knowledge from Molecular Dynamics simulations has broad implications

There has been an exponential growth of long-time scale MD simulations of proteins and protein complexes, providing rich information on their dynamic properties (6,34–41). However, there is a need to shift the usual paradigm of doing selective analysis of MD simulation trajectories to a more data-driven analysis using the interpretable network-centered methods established in the Computational Systems Biology field for large-scale multimodal data analysis (42–46). The existing MD simulations secondary data analysis approaches often employ dimensionality reduction techniques, primarily principal component analysis (PCA), which suffer from the inherently low interpretability, and from the related issue of the inability to separate induced and direct (non-transitive) dependencies (47–49). Residue-based network models have been used to study the contact relationships in static structures (47–50), but these network models are residue-centric and do not bring out the correlations among inter-residue contacts that are distant or allosteric (50). Importantly, there are currently no network-centric systems biology tools that represent residue pairs as network nodes for inferring dependencies (or cooperativity) between inter-residue contacts from dynamics data. Doing so would harness the rich statistics of heterogeneous molecular interactions embedded within long time-scale MD trajectories for predicting cooperative interactions and identifying hotspot contacts not discernable through conventional analysis.

Our goal is to use an interpretable unsupervised machine learning network-centered methodology, BN modeling, with the protein-protein interface (GPCR:G protein interface in this context) contacts as the nodes in the network model. We previously established BNOmics as a robust and scalable BN platform for analyzing large scale multimodal biomedical datasets (42). We have now applied BNOmics to MD simulation data, utilizing the capacity of BNs to infer nonlinear non-transitive dependencies (suggesting directional causality) among different parts of the GPCR:G protein interface (32,33). BN modeling has been used with the goal to decipher the direct dependencies and the cooperative behavior of the inter-residue contacts between GPCR and G proteins during the MD simulations, thus pinpointing the residue pairs that contribute significantly to G protein coupling. We have chosen BNs as the primary network-centric method for this study because the alternatives suffer from low interpretability (dimensionality reduction, e.g., PCA), do not filter out spurious (induced, or transitive) dependencies (correlation networks (51,52)) or rely on the residues as the input (Gaussian networks (53–55)).

## Results

### BN modeling methodology as applied to the GPCR:G protein interface residue contacts data

The workflow developed in this study to apply BN modeling to MD simulation trajectories is shown in Figure 1. A BN is a graphical representation of the joint probability distribution of a set of random variables (nodes in the network), where the network topology (directed acyclic graph) reflects relationships among the nodes (i.e., which nodes are directly dependent on, or conditionally independent of, other nodes). In this application the nodes are the GPCR:G protein interface residue contacts mapped using MD simulations of the GPCR:G protein complexes in explicit lipid bilayer. The residue contacts are binary (1 or 0, contact present or absent) as shown in the middle panel of Fig. 1. Edges connecting the nodes indicate direct (non-transitive) dependencies, with no linearity assumptions. In our case, the edges imply non-parametric (multinomial) statistical correlation between the protein residue pairwise contacts. In other words, the edge between the two contacts indicates that they co-evolve in a concerted manner during dynamics. While edge directionality in the BN might suggest causation, what is more important is that the directed BNs eschew spurious dependencies induced by multicollinearities (32,33,42). Each edge in the BN is accompanied by its edge strength, which is proportional to the ratio of the marginal maximum likelihood of the network with the edge to the one without, given the data (46). Looking at the BN edge interpretation from the biological perspective – while we know that the three-dimensional static structures show certain GPCR:G protein residue contacts, the information on which of these residue contacts contribute cooperatively to the G protein coupling strength and selectivity is not evident from enthalpy and temporal persistence alone. The BN edge strength analysis using the GPCR:G protein residue contacts as nodes will provide a quantitative estimate of the co-operativity among the pairwise interactions at the interface. The sparse nature of the directed BNs will prioritize the non-transitive, direct, dependencies between the residue contacts. Deconvolution of the contact frequency patterns using BN discriminates true cooperativity hotspots that mechanistically drive the propensity of other contacts at the interface, as opposed to contact correlations that are mere co-occurrences. Comparison of BNs across multiple G protein coupled complexes can be used to identify common and unique residue hotspots that are involved in G protein selectivity and promiscuity.

**Figure 1.**
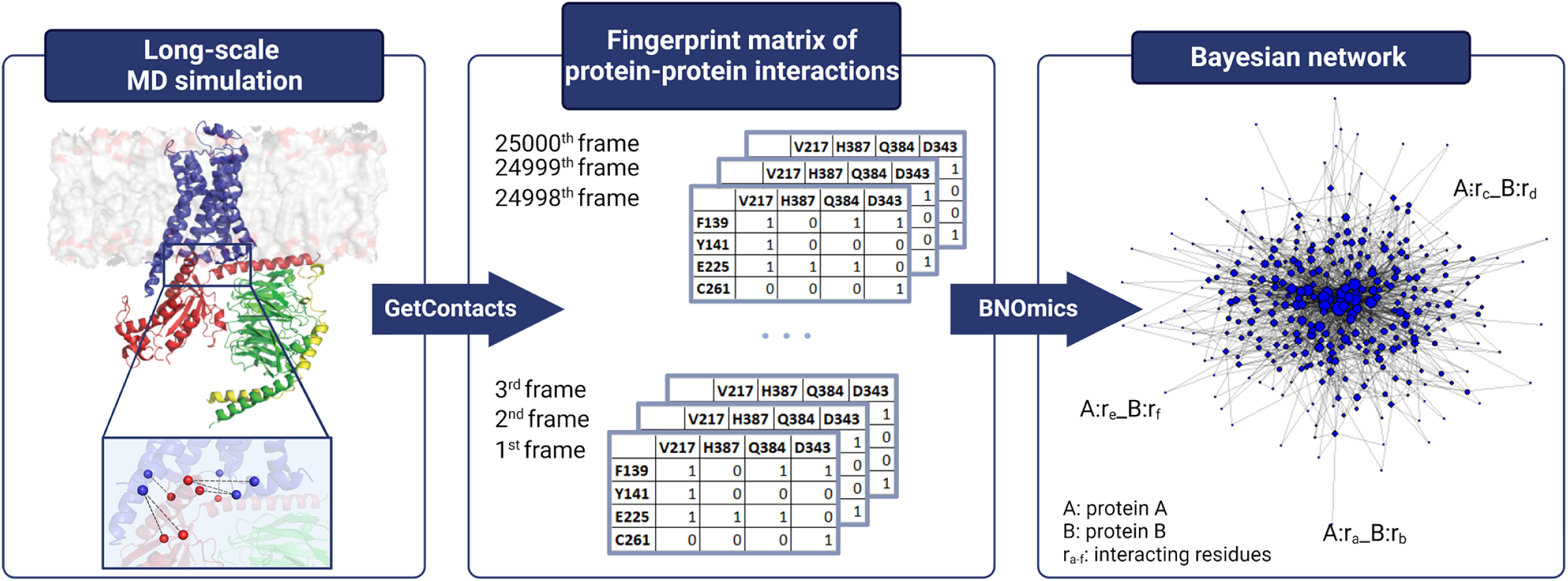
**Workflow combining MD simulations and Bayesian network (BN) models to delineate the GPCR:G protein interface residue contacts that show cooperativity for G protein coupling**. The left panel shows the GPCR:G protein interface residue pairs that display direct interaction as calculated from the MD simulation trajectories of each of the GPCR:G protein system using the program “GetContacts” (73). The middle panel shows the stack of fingerprint matrices for the GPCR:G protein residue contacts derived for every MD snapshot. The right panel shows the BN model derived for each GPCR:G protein complex using BNOmics software (42).

### MD simulations show expanded repertoire of GPCR:G protein contacts

When we started this study there were three dimensional structures available for 40 Gi-coupled receptors, 40 Gs-coupled receptors, and 11 Gq-coupled receptors. However, 9 out of 11 structures of the Gq-coupled receptors contain chimeric Gq protein with only the last 5 to 10 amino acids of the C-terminus of the G protein being that of Gq. Since such structures may not reflect how a wild type Gq would couple to GPCRs, we took only the two GPCR:Gq structures that contained the wild type Gq protein. Therefore, we performed MD simulations on 2 Gs (β_2_AR: 3SN6 (56); A_2A_R: 6D9H (57)), 2 Gq (H_1_R: 7DFL (58), AT_1_R: 7F6G (59)) and 2 Gi coupled (A_1_R: 6GDG (60,61), 5HT_1B_R: 6G79 (61)) receptors. We performed multiple runs of all-atom MD simulations in explicit POPC bilayer with cholesterol (see Methods) with an aggregate simulation time of 5μs for each GPCR:G protein system. We analyzed the number and nature of GPCR:G protein interface residue contacts in each complex in the static three-dimensional structures and in their respective MD simulations trajectories (Fig. 2a). To enable comparison across multiple GPCR:G protein interface contacts, we used the GPCR residue numbering system (62) and Common G protein numbering (63).

**Figure 2.**
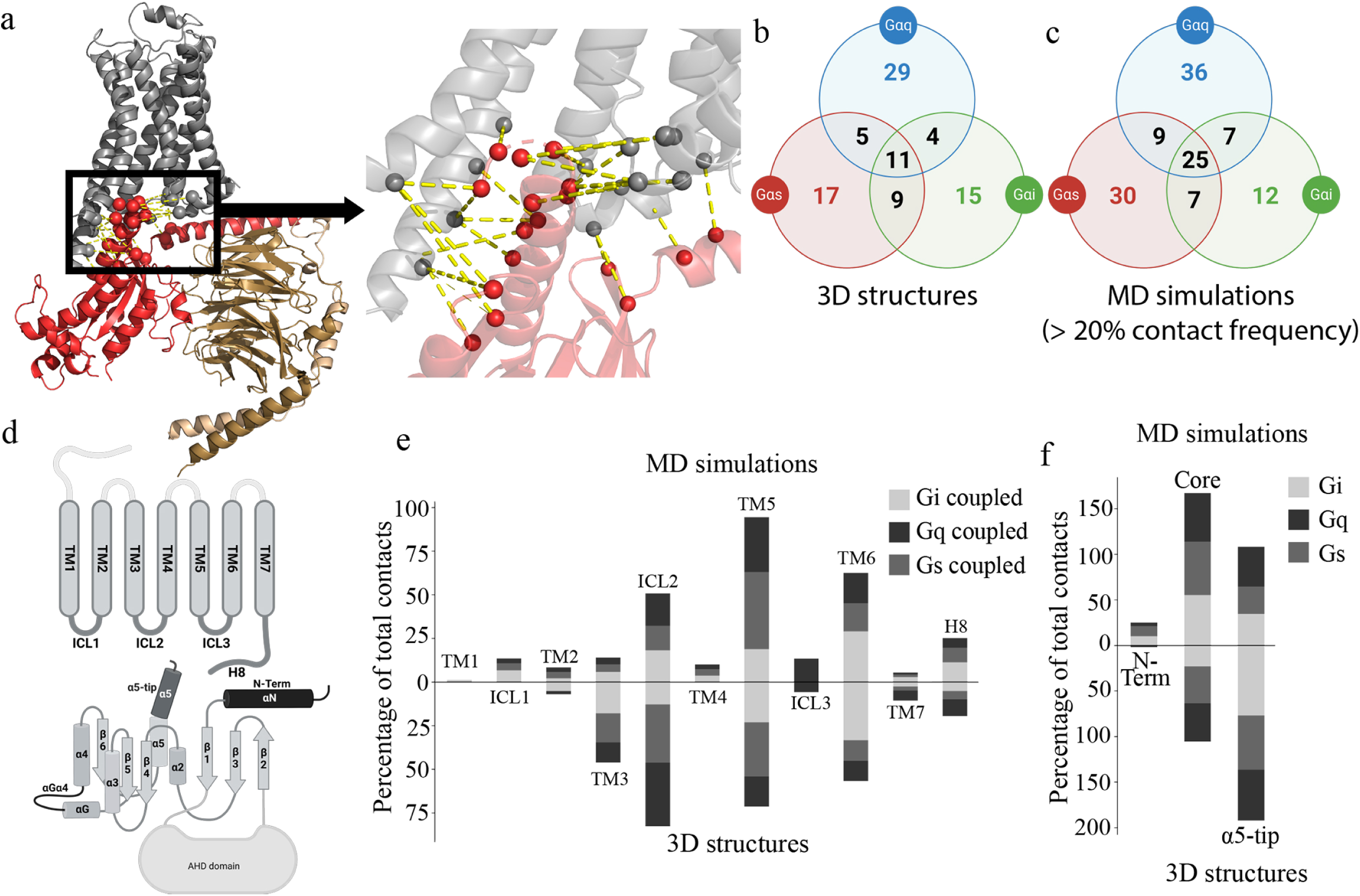
**MD simulations expand the repertoire of GPCR:G protein interface interactions**. **a.** General representation of GPCR:G protein complex. Area highlighted by black rectangular is the GPCR:G protein interface that was used in further analysis. **b**. Venn diagram of GPCR:G protein residue contacts present in the 3D structures (green circle – Gi-coupled receptors: 6D9H, 6G79; red circle – Gs-coupled receptors:3SN6, 6GDG; blue circle – Gq-coupled receptor: 7DFL, 7F6G). **c**. Venn diagram of persistent GPCR:G protein residue contacts formed during the MD simulation. **d**. GPCR and G protein schematic. **e**. Bar plot demonstrating how residue contacts are distributed over GPCR structural elements in the CS and during MD simulation. (62). **f**. stacked plot of the percentage of GPCR:G protein residue interactions observed in the 3D structures and in MD simulations located in different structural regions of the G protein subtypes.

We first compared the common and distinct contacts observed in the three-dimensional structures of GPCRs bound to the three G protein subtypes with those identified from the MD simulations (Fig. 2b,c). As shown in Fig. 2b, the three-dimensional structures of the 6 GPCR:G protein complexes showed 39 GPCR:G protein contacts combined in the two Gi coupled receptors, 42 contacts combined in the two Gs coupled receptors and 49 contacts combined in the two Gq coupled receptors. These contacts were formed by 20-25 distinct residues on the GPCR, and 14-22 on the G protein. We then did further comparison by classifying these contacts by structural regions in which they are located (Fig. 2d-f). As illustrated in Fig. 2d, we identified the GPCR interface residues as located in transmembrane (TM) helices and intracellular (IC) loops (see Methods). G protein was divided into three domains, N terminus (first 29-36 residues), tip of the α5 helix (last 10 amino acids at the C terminus) that inserts into the GPCR IC region, along with the rest of the structure broadly designated as the core region (Fig. 2d). TM3, TM5, TM6 and ICL2 contain the greatest number of contacts, although the distribution of contacts among these domains differed in the G protein subtype. For example, 33% of the total contacts are from residues in TM6 in Gi coupled receptors compared to 12 % in Gs and Gq coupled receptors (Fig. 2e). In contrast, the contribution of residues in ICL2 is 33% and 36%, respectively in Gs and Gq coupled complexes compared to 13% in Gi coupled complexes. Residues on TM5 make more contacts in Gi and Gs coupled receptors compared to Gq coupled receptors. It should be noted that many of these static structures are missing multiple stretches of residues in ICL3 and hence not included in the MD simulations. Therefore, the contribution by ICL3, while previously shown to be important for GPCR:G protein coupling and selectivity (64), is not included in this study (Fig.S2a).

On the G protein side, 77% of total interactions are in the α5-tip, compared to ∼60% of total residue contacts in the Gs and Gq proteins (Fig. 2f). The number of GPCR:G protein interactions with residues in the G protein core region expands during MD simulations compared to those in the static three-dimensional structures. In contrast, the number of contacts made by residues in the α5-tip decreases during the MD simulations. There are 25 common contacts among all G protein subtypes that come mainly from ICL2 and TM6 in the GPCR and from the α5-tip of the G protein among all the three G protein subtypes (Fig. 2c). The G protein N terminus mediated interactions were formed during MD simulations in all G protein subtypes, mainly contacting ICL1, ICL2, and TM4 residues in the GPCRs. Notably, the N terminus mediated interactions were missing in all the static structures. As shown by Jelinek *et al* (65), using chimeric G protein constructs (discussed in detail in the last section of the results), N terminal G protein residues contribute towards G protein selectivity. Hence, the new contacts observed during MD have functional significance, as confirmed by experimental studies. The increase in the number of interface contacts in the MD simulations comes from newer interactions being formed with the neighboring residues during the dynamics, specifically from the intracellular end of TM5 (Fig. 2e). Each G protein residue contact with multiple GPCR residues or vice versa. For example, residue 34×51 in the three-dimensional structure of adenosine A_1_R receptor contacts two G protein residues (H5.12 and H5.15) while during the MD simulations the number of contacts increases to 4 persistent contacts (greater than 20% frequency in the MD simulations) made with residues H5.08, H5.12, H5.15 and S3.01. These persistent contacts made by the GPCR:G protein interactions in the MD simulations are provided in the Supplementary Data S1. In summary, the analysis of GPCR:G protein contacts, based solely on static structures, may give an incomplete picture of the interaction interface, since many contacts, that are observed during dynamics, would be missed.

The differences in the distribution of residue contacts among the different G protein subtypes also changed significantly from the static structures during MD simulations (Fig. 2d, e). For example, the ICL2 contribution among the Gi coupled receptors was the smallest among the three G protein subtypes, according to the static structures. During MD simulations, the ICL2 contribution in Gi increased to match that of Gq (18%), while its involvement in Gs reduced to 14% (Fig. 2e). Residues on the TM5 contribute significantly in the Gs coupled receptors mainly coming from the longer hgh4-loop in Gs that makes interactions with the intracellular end of TM5 and N-terminus of the ICL3. These interactions have been shown previously to be crucial for receptor-Gs protein coupling(64) (Fig. S2b). Moreover, there are regions that are specific to only particular subtypes, such as residues on the intracellular side of TM1 that make contacts in the Gi and Gq coupled receptors but not in the Gs-coupled receptors (Fig. 2e). Also, only Gq coupled receptors show ICL3 mediated contacts in our analysis. However, this is attributed to the fact that only one of the six structures studied here has the full ICL3 resolved (Gq coupled angiotensin II receptor 1 AT_1_R). Since the ICL3 is not resolved in other structures we will not be counting these contacts in our analysis. In summary, the number of GPCR:G protein interactions increase during the MD simulations showing that the G protein core is equally involved in interacting with the GPCR as the α5-tip of the C-terminus.

### Bayesian network models show distinct GPCR:G protein interaction residues with high cooperativity

Given the increased repertoire of GPCR:G protein interactions, we wanted to identify the interactions that show high level of cooperativity using a network model. Using the GPCR:G protein residue contact fingerprint matrix generated for each MD simulation snapshot (Supplementary Data S2-S7), shown in the middle panel of Fig. 1, we derived the BN models for each system using the BNOmics (42) software as described in the Methods section. A BN was derived for each GPCR:G protein complex, with each node in the network representing a GPCR:G protein residue contact. Evaluation of the robustness of the network models was conducted as described in the Methods section. Random local perturbations in the topology of the network revealed that the scores of the original models remained the highest, suggesting that the topology found by BNOmics is close to the globally optimal topology given the data (Figure S3a). This was not surprising, given the favorable dimensionality of the data. In addition, we tested the robustness of our models to sampling by conducting a series of sensitivity experiments. As expected, all scores resided in the 0.99 confidence interval, implying that our models were not sensitive to the sampling schedule variations (Fig. S3b).

To identify the nodes that are centrally connected in the BN (Fig. 3a), we used the network property “edge strength”, which is a measure of variable dependency, proportional to the ratio of the marginal maximum likelihood of the network with the edge to the one without, given the data (46). The total node strength, also defined as “cooperativity” of each node, was calculated as a sum of the edge strengths of its connections (edges) (Fig. 3b). Since we use residue pairs as nodes in this network model, the total edge strength for each node may be treated as a quantitative measure of the cooperativity with other nodes in the network. This is because BN analysis can identify co-dependencies between the variables in the dataset which translate to cooperative movements in the dynamic protein systems. The cooperativity ranking of all residue contacts of each GPCR:G protein system is provided in the Supplementary Data S8. As an example, Fig. 3b demonstrates how node strength of each residue pair differs depending on the number and strength of connections it makes in the BN model. A close-up view of a section of the network is shown in Fig. 3b, where the central node 7×55_G.H5.24 is the most cooperative interaction in the β_2_AR-Gs complex. Compared to other nodes, this node has a notably higher node strength due to the strength and number of its connections, thus making it crucial for the network structure (Fig. 3b). We used the calculated cooperativity (Fig. 3b) of each node to assess its importance in GPCR:G protein coupling.

**Figure 3.**
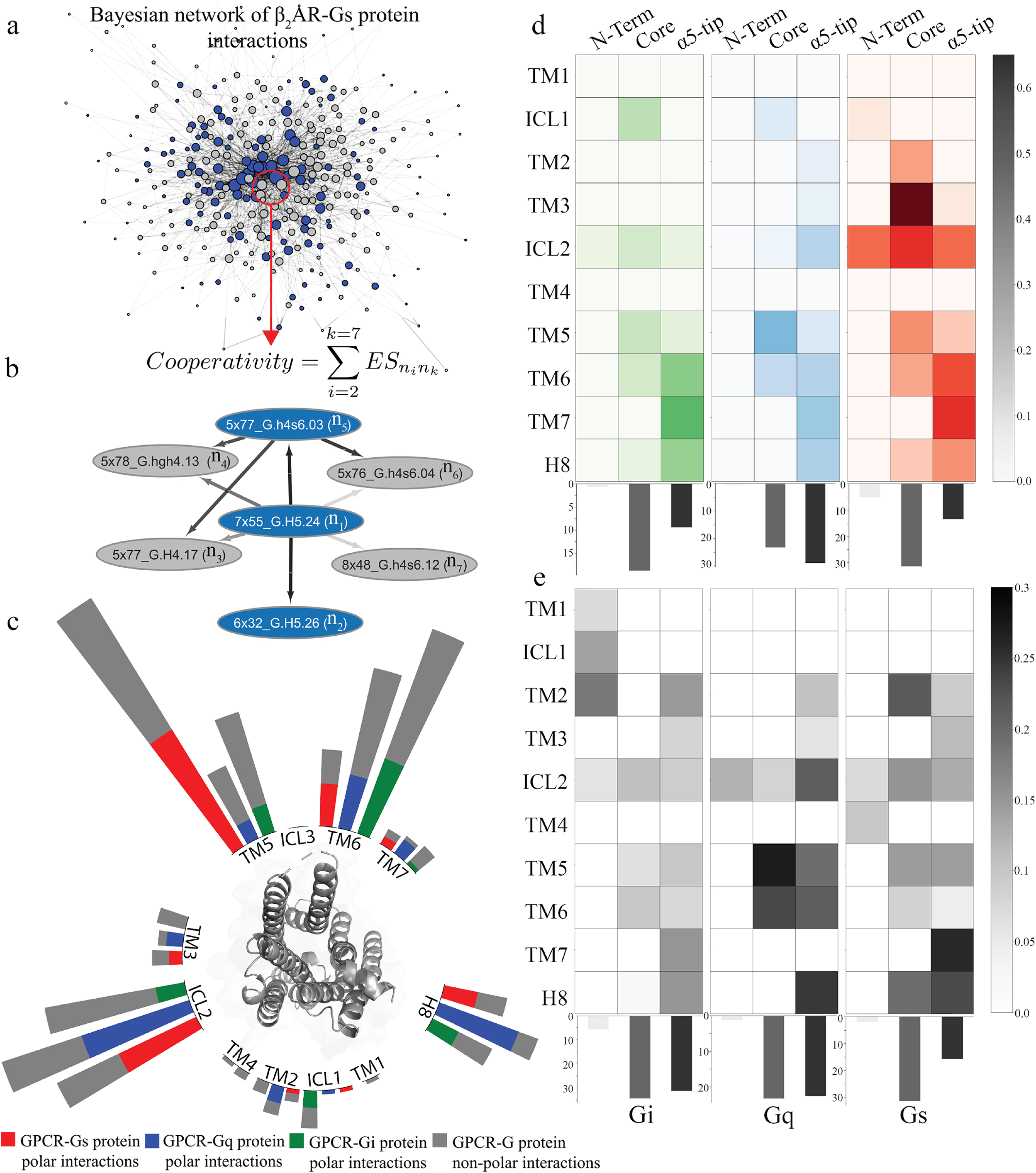
BN modeling-based analysis of GPCR:G Protein interaction residues and their cooperative dynamics in different G Protein subtypes. a. A complete Bayesian network model of the GPCR:G protein interaction residues in β2 adrenergic receptor: Gs complex. Arrow thickness corresponds to the edge weight. Blue node – polar interaction, grey – non-polar interaction. **b.** Formula used for calculating the cooperativity of each node in the BN. An illustrative subnetwork from the β_2_AR-Gs complex centered on the most cooperative node 7×55_G.H5.24. **c.** Radial plot of cooperative interactions and their distribution over GPCR structural elements. Blue – polar interactions in Gq coupled receptors, red – polar interaction in Gs coupled receptors, green – polar interactions in Gi coupled receptors. Grey-hydrophobic interactions in the all the G protein subtypes. **d.** Heatmap of the fraction of polar cooperative interactions each of the G protein structural region forms with each TM helix of the GPCR. Each TM-G protein region fraction was corrected on the number of total interactions made by this G protein region, to account for differences in the sizes of G protein structures. The darker the color, the larger the number of cooperative nodes in the structural region of GPCRs and G proteins. Left to right: Gi coupled, Gq coupled, Gs coupled. Bar plots are showing the percentage of cooperative interactions formed by the different G protein structural regions. **e.** Heatmap of the fraction of non-polar cooperative interactions each of the G protein structural region forms with each TM helix of the GPCR. Each TM-G protein region fraction was corrected on the number of total interactions made by this G protein region, to account for differences in the sizes of G protein structures. The darker the color, the larger the number of cooperative nodes in the structural region of GPCRs and G proteins. Left to right: Gi coupled, Gq coupled, Gs coupled. Bar plots are showing the percentage of cooperative interactions formed by the different G protein structural regions.

We sorted the nodes by their cooperativity score (defined in Fig. 3b) and analyzed the structural regions of the GPCRs and the G proteins in which the nodes (the most cooperative ones) in the top quartile in each complex are located. We categorized the nodes (the type of GPCR:G protein interactions) into polar and non-polar interactions. As shown in Fig. 3c, TM5, TM6, ICL2 and helix 8 (H8) contain the most cooperative nodes in the network for all the three G protein subtypes. On the G protein side, as seen in Figs. 3d and 3e, the percentage of the cooperative nodes in the G protein core and α5-tip are similar for the Gq (47% and 52%) and Gi (53% and 40%, respectively) proteins. The core region shows a significantly higher percentage (63% and 30% respectively) of cooperative interactions in the Gs protein. This suggests that both the core and alpha5-tip, despite the latter being more extensively studied (4,6,7,13,22,66–69), play an equally important role in cooperativity. This is a significant finding, challenging the conventional understanding and emphasizing the need to focus on the core region as an integral part of the GPCR signaling mechanism. Additionally, cooperative nodes are as often non-polar as polar interactions showing that non-polar contacts play an equally important role in the GPCR:G protein interface.

Some of the Gi protein specific cooperative non-polar contacts regions are in the N terminus of the G protein forming contacts with TM1, TM2 and ICL1. These interactions are not present in the Gs and Gq protein coupled receptors. Although the number of contacts formed by residues in TM4 (Fig. 2d) is the same across all the G protein subtypes (Fig. 2d), the high scoring cooperative nodes of TM4 are appearing only in Gs and Gq coupled receptors. Taken together these results show that BN models from GPCR:G protein contact fingerprints can be used to study the cooperativity of the GPCR:G protein complex interactions. We identified key cooperative GPCR:G protein contacts via node strength analysis and revealed that regions TM5, TM6, TM7, and ICL2 contain the most cooperative nodes, with different balance of polar and non-polar interactions among G protein subtypes.

### Bayesian network models show co-operativity among distant (allosteric) residue pairs in the GPCR:G protein interface

BN modeling offers a valuable approach for analyzing the dependencies between variables. In the BN models where the nodes are GPCR:G protein contacts, many distant nodes showed strong dependencies, indicating cooperative allostery across the interface. In the BN model, edges connecting the nodes are established in the probability space, rather than by spatial distance, since BN is agnostic of the structural coordinates of the protein atoms. While it is relatively easy to study the influence and co-dependency of closely located residues or contacts using structure-based network models, understanding the cross-communication between structurally distant regions during protein-protein complex dynamics poses a challenge. To investigate these long-distance dependencies and the effectiveness of BN models in capturing them, we classified the edges in the BN models into two categories: neighboring dependencies (Cα atoms of GPCR and G protein residues within 10Å proximity) and allosteric dependencies (Cα atoms of GPCR and G protein residues more than 10Å apart). We then compared the proportions of allosteric vs neighboring edges within the Markov neighborhoods, the set of nodes that are directly connected to node of interest, of top nodes (having the most BN connections) of the three G protein subtypes (Fig. 4a). We found that there are multiple allosteric edges connecting each of the top scoring cooperative nodes across all three G proteins, with varying edge strengths. Our observations indicate that the dynamics of GPCR:G protein contacts are not only influenced by their local spatial neighborhood, but by contacts that are formed far off in the interface, thus establishing a profound contribution of allosteric communication across the G protein structures.

**Figure 4.**
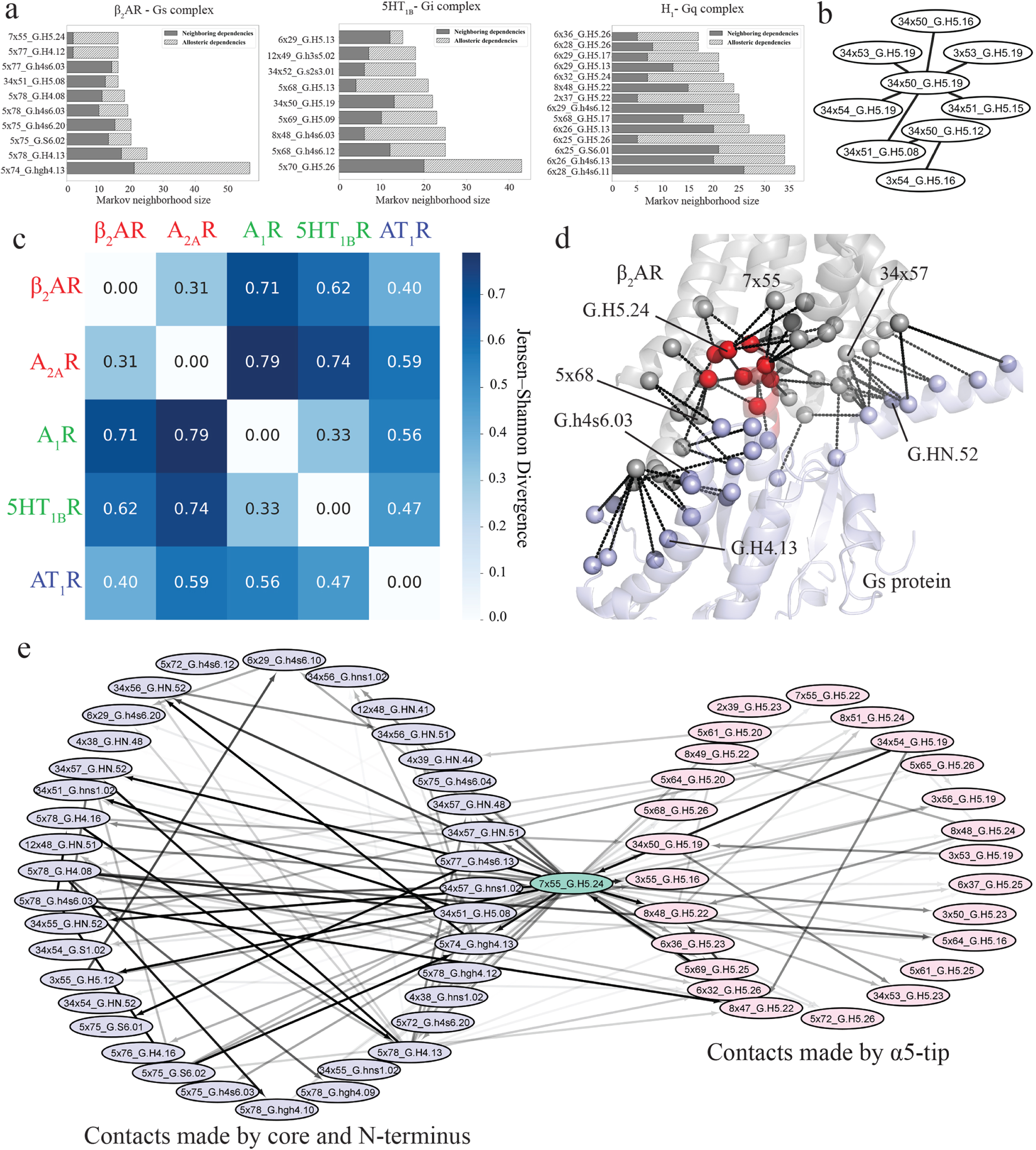
Comparative Analysis of Markov Neighborhoods in GPCR:G Protein Complexes: Highlighting Core Interactions and Distinctive Contact Regions. a. The Markov neighborhoods of the most connected nodes of Gi, Gq and Gs coupled GPCR complexes (left to right). Only the neighborhoods larger than 10 nodes were displayed. **b.** The largest common subnetwork of GPCR:G protein interactions that appears at least in 5 out of 6 moralized versions of directed BN models. **c.** The heatmap representation of the JSD matrix of joint probabilities given the topology of the largest common subgraph. Red color represents Gs coupled receptors, green – Gi coupled, blue – Gq coupled. H_1_R receptor was omitted due to the absence of this common subgraph in the moralized graph. **d.** The structural representation of the Markov neighborhood of 7×55_G.H5.24. Cα atoms of residues, making contacts, are represented as spheres. The receptor is represented by grey color, the core of G protein – blue, the tip – red. **e.** The Markov neighborhood of the most connected node, 7×55_G.H5.24 (light-green), of the β_2_AR-Gs complex. Light-red color corresponds to the contacts made by the α5 tip, light-blue color – contacts made by the core of the G protein. The color intensities of the edges are proportional to the edge strength.

To analyze the similarities among the BN models of different G protein subtypes, we have performed moralization of the directed graphs (see Methods) obtained from BNOmics to enable direct comparison of dependencies across the networks of each system. During moralization, we converted the directed network architecture of the BN to an undirected one by adding a new edge to every pair of “parental” nodes that showed dependency towards a third common “offspring” node (70). Doing so allowed us to compare network topologies across multiple BNs. Next, we identified the largest common subnetwork of nodes, i.e., subnetwork that appear in the BN models of all the G protein subtypes, as demonstrated in Fig. 4c. This subnetwork is almost exclusively formed by the contacts of ICL2 with α5-tip, indicating that interaction between these two structural regions is important for coupling to all G protein subtypes. Upon obtaining this subnetwork, we evaluated the joint probability for each complex for all nodes. We then estimated the divergence between each joint probability using Jensen-Shannon Divergence (JSD) with the results reported in Fig. 4b. We observed that the distance between complexes of the same subtype is relatively small, e.g., β_2_AR and A_2A_R, which means that probabilistic relationships of the nodes in subgraph are similar for these two systems. However, comparing joint probabilities of GPCRs coupling to different G protein subtypes, e.g., β_2_AR and A_1_R, reveals that there is a significant difference between their probabilistic relationships despite the same topology. This means that even though the subset of interactions between ICL2 and α5-tip is observed across all G protein subtypes, the relationship between contacts inside this subset is G protein dependent. In particular, the residues that formed the contacts in this common graph have been shown to be important for GPCR and G protein coupling selectivity (5,6,66,71,72). Interestingly, the common subgraph in Fig. 4b does not include any G protein residues from the core or N-terminus regions. The pattern of cooperativity involving these regions is more specific to G protein subtype and may be strongly involved in determining selectivity.

To demonstrate the significance of cooperative allosteric crosstalk across the GPCR:G protein interface for G protein coupling (Fig. 4d), the network, involving the most connected node, 7×55_G.H5.24, in the β_2_AR-Gs complex system, is showcased in Fig. 4e. Even though the most central node is made by a residue in the α5-tip (H5.24), many of the “allosteric nodes” in its neighborhood are formed by the residues in the core and the N-terminus of the G protein. For example, the strong edges between 7×55_G.H5.24 and 12×48_G.H4.08, 5×78_G.hgh4.09, 34×56_G.hns1.02 clearly demonstrate the allosteric cooperativity of the node 7×55_G.H5.24 Mapping the Markov neighborhood of 7×55_H5.24 to the β_2_AR-Gs three-dimensional structure (PDB: 3SN6 (56)) shows that the cooperative dependencies of this contact span the entire GPCR:G protein interface (Fig. 4d). We propose that such cooperative crosstalk allows multiple contacts to be formed simultaneously across the entire G protein interface, when the G protein couples with the receptor. This ensures sustained and robust coupling between the two proteins, while in the absence of cooperative interactions, such coupling would be transient. We further posit that such mechanisms are broadly applicable to protein-protein coupling. Here we also demonstrate that these cooperative contacts can be identified for any protein complex using BN models combined with MD simulations.

### High cooperativity GPCR:G protein contacts are crucial for the selectivity of G protein coupling

Uncovering the residues in the GPCR:G protein interface that confer G protein coupling selectivity is critically important to designing selective drugs. Ample number of prior studies have highlighted the importance of residues in the C-terminus tip of the G protein to be important in G protein coupling selectivity (2–4,15,66,73). Although the core region of the G protein has also been shown to be important in G protein selectivity (4), the specific residues in the core region that are important for selectivity are not known. We hypothesized that the cooperative nodes identified through the BN approach play an important role in conferring G protein selectivity and sought validation using experimental observations.

To assess the significance of key BN nodes in selective G protein coupling, we prioritized the G protein residues that are in the top quartile of cooperativity score in any pair of G proteins (Gs, Gi and Gq). We also used a second criterion to choose highly cooperative residues in the G protein that are not conserved across the G protein subtypes (Fig. S4a). We calculated the cooperativity score (mean node strength of unconserved residues in the top quartile of cooperativity score in the structural region of interest) for each structural region of the G protein (Fig. 5a). As seen in Fig. 5b, we found that the α5-tip and the rest of the α5 helix called the α5-core of the C-terminus of the G protein are the highest scoring regions, with the α5-tip to be the top contributor of cooperativity. Interestingly, we also found residues with high cooperativity scores in several other regions of the G protein core, notably the β2β3 loop and the β6 segment. To validate these predictions on the selectivity determinants in the G protein core we compared them to the experimental data. Jelinek et al. generated chimeras by substituting different regions at the GPCR:G protein interface in the Go structure with the corresponding amino acid sequences from Gq (65). Using FRET based assays, the coupling efficiencies (EC50) of these chimeras were measured for two GPCRs, the muscarinic M_3_ (M_3_R) and histamine H_1_ receptors (H_1_R), both of which show stronger affinity for Gq compared to Go. By analyzing the change in EC50 of the chimeras over wild type Go, the authors identified several key hotspot regions in the G protein structure that significantly contribute to the selectivity of M_3_R and H_1_R towards Gq (red stars in Fig. 5a). Besides the α5-core (first 10 residues of the C-terminus, Fig. 5a) and α5-tip (last 11 residues, α5-tip), the other regions reported as hotspots include a hinge region between the N terminus and β1 segment (N-term) and the loop connecting β2 with β3 (β2β3). The β2β3 loop was reported as one of the highest selectivity hotspots by Jelinek et al. Plots of calculated cooperativity score versus measured changes in EC50 for the chimeras relative to the wild type Go (ΔlogEC50) for muscarinic acetyl choline receptor M1R and histamine receptor 1, H1R as reported by Jelinek et al (65) are shown in Figs. 5c, d. For both receptors, the cooperativity scores showed an inverse correlation with ΔlogEC50, indicating a non-trivial connection between BN node strengths and their contributions towards G protein selectivity. This shows that BN analysis of MD simulations data can be used to predict the selectivity determinants of the GPCR:G protein coupling. We also found additional cooperativity hotspots predicted by BN that were not reported previously, e.g., the β6 segment. Future advancements of this approach will involve harnessing the information from BN subnetworks in designing selectivity modulating mutations or designing peptides or small molecules targeting G proteins and GPCR:G protein interfaces.

**Figure 5.**
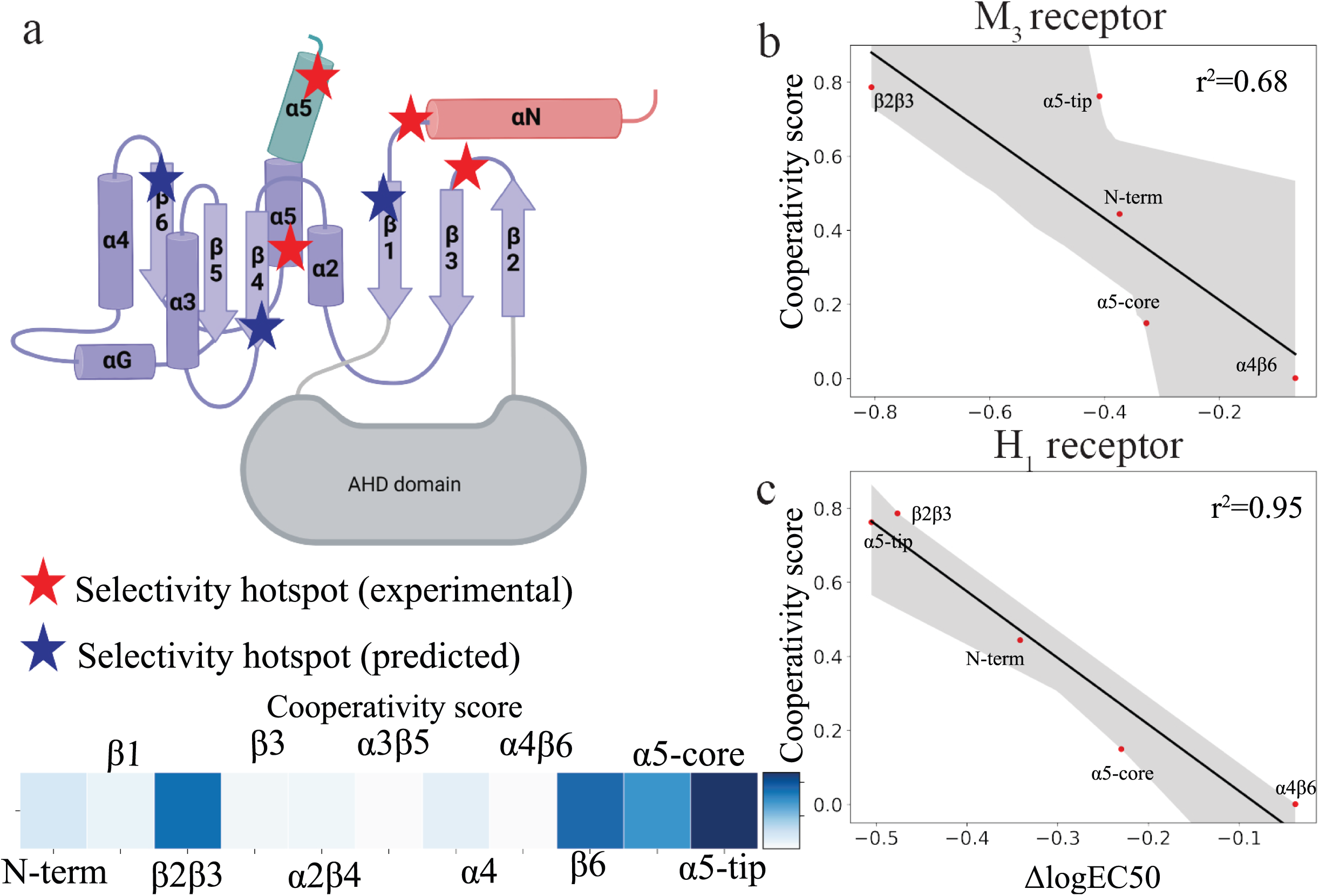
Comparative Analysis of G Protein Selectivity Hotspots: Experimental vs. BN Model Predictions. a. Schematic representation of G protein secondary structure. Red stars represent regions identified as selectivity hotspots by Jelinek et al (65), blue stars represent electivity hotspots predicted by BN models. Heatmap represents the cooperativity score of proposed selectivity hotspot based on the BN model of H_1_-Gq complex. **b.** ΔlogEC50 values of measuring of M_3_-Goq chimera complex coupling from Jelinek et al. plotted against node strength of predicted selectivity hotspots by BN modeling. **c.** ΔlogEC50 values of measuring of H_1_-Goq chimera complex coupling from Jelinek et al. plotted against node strength of selectivity hotspots identified by BN modeling.

## Discussion

Although it is known that cooperative interactions play a critical role in protein-protein interactions, there is a serious lack of methods to uncover these interactions. In this study, we have adapted Bayesian network modeling that has been used extensively in generalist multimodal network construction and dissection, to MD simulation trajectories to analyze the cooperativity among the pairwise GPCR:G protein interactions. The MD simulations starting from the three-dimensional structures revealed multiple additional interactions that were absent in the static structures, thereby enriching the comparison among different G protein subtypes. Our analysis revealed specific regions of GPCRs and G proteins that play critical roles in mediating the interactions between these proteins. Residues on intracellular loop 2 (ICL2), transmembrane helix 5 (TM5), and TM6 consistently formed contacts with the G protein in all studied complexes. This observation is in line with previous studies (5,17,18,23–25,56,68,74) highlighting the importance of these regions in GPCR:G protein coupling. However, our analysis revealed significant differences in contact frequencies involving these regions among the different G protein subtypes, indicating that such differences may contribute to G protein specific coupling preferences.

BN analysis of MD simulation data revealed the GPCR:G protein contacts that are highly cooperative, as indicated by the size of their Markov neighborhoods. These interactions were primarily located on ICL2, TM5, and TM6 of GPCRs and in the core and C terminal regions of the G proteins. Notably, we found a cooperative contact subnetwork formed by residues in the ICL2 and the α5 helix of the G proteins that were common in the BN of majority of the systems studied here. Such common cooperative hotspots may be required for the recognition of G proteins by GPCRs irrespective of the G protein subtype. We also found multiple cooperative contacts involving the core and N terminus of G proteins that are specific to the different subtypes. These contacts may confer selectivity to the GPCR:G protein recognition through further interface stabilization.

One important finding in this study is that the mutually cooperative contacts at the G protein interfaces are not always spatially close, thereby suggesting that cooperativity at protein-protein interacting interfaces may not be strictly mediated by structurally close contacts. A large fraction of the probabilistic Markov neighborhoods of highly cooperative contacts are formed by distant contacts, thereby indicating that allostery plays a crucial role in conferring cooperativity within protein-protein interface. We found that these cooperative allosteric contacts are distributed throughout the G protein interface suggesting that the entire interface acts synergistically in stabilizing the coupling with GPCRs. Finally, the cooperativity scores of individual G protein regions correlated well with the changes in GPCR coupling efficiencies obtained from chimeric G protein studies. This suggests that our computed cooperativity measures could be used to prioritize specific G protein regions for designing mutations and targeted therapeutics to modulate the selectivity of G proteins towards their cognate GPCRs.

In conclusion, our study advances the understanding of GPCR:G protein interactions by combining network-based analysis and MD simulations. The results highlight the cooperative nature of these interactions and identify key residues and regions involved in GPCR:G protein coupling. The findings provide valuable insights into the dynamic behavior and selectivity of GPCR:G protein signaling, opening avenues for the design of selective drugs targeting specific G protein subtypes. Further experimental investigations based on the predicted residues and network models will contribute to a deeper understanding of the intricate mechanisms governing GPCR signaling and its therapeutic implications.

## Methods

### Preparation of Ligand and Receptor Model

All six GPCR:G protein complexes were prepared for MD simulations using corresponding EM structures (A_2A_R with mini-Gs:6GDG, A_1_R with Gi2:6D9H, 5-HT_1B_R with Go: 6G79, AT_1_R with Gq: 7F6G, H_1_R with Gq: 7DFL) or X-ray structures (β_2_AR with Gs: 3SN6). All mutations in receptors and G proteins were reverted to wild type residues using Maestro (Schrödinger Release 2020-1: Maestro, Schrödinger, LLC, New York, NY, 2020.). All ligands were parameterized by ParaChem (https://cgenff.umaryland.edu, (75)). Missing sidechain residues and loops, each having fewer than 5 absent amino acids, were integrated into the 3D structures of the GPCR:G protein complexes. Residues within 5 Å of mutation sites were minimized using MacroModel, with all backbone atoms subjected to position restrictions. The protein chain termini were capped with neutral acetyl and methylamide groups, and the histidine protonation states were assigned using Maestro protein preparation wizard. The complex was then placed into an explicit POPC bilayer membrane using the PPM 2.0 function of the Orientation of Proteins in Membranes (OPM) tool (76), and hydrated with TIP3 water containing 0.15 M NaCl in CHARMM-GUI (77,78). The final simulation system dimensions were approximately 125 Å × 125 Å × 170 Å. All of the simulation systems were characterized by CHARMM36m force field (79).

### MD Simulations for GPCR Complexes

GROMACS2019 package (80,81) was used to conduct all MD simulations using 2 fs integration time step. The simulation systems, once prepared, underwent an initial minimization with position restraints of 10 kcal/mol·Å2 on all heavy atoms encompassing the GPCR, G protein, ligand, and lipids. This was succeeded by a 1-ns heating phase that escalated the temperature from 0 K to 310 K under the NVT ensemble, utilizing the Nosé-Hoover thermostat. Subsequent to this, the system was subjected to an equilibration simulation within the NPT ensemble. Here, the initial 1 ns had the aforementioned 10 kcal/mol·Å^2^ position restraint. This restraint was then progressively reduced: first to 5 kcal/mol·Å^2^ and then diminishing to 1 kcal/mol·Å^2^ in decrements of 1 kcal/mol·Å^2^. Each decrement was accompanied by a simulation spanning 5 ns. The final phase of the equilibration comprised a 50 ns simulation, executed devoid of any position restraints. The final snapshot of the equilibration step served as the initial conformation for five production runs (each last for 1 μs), which were initiated with randomly generated velocities. The simulation pressure was coupled to a 1 bar pressure bath and controlled using the Parrinello-Rahman method (82). Nonbond interactions had a cutoff distance of 12 Å, and the Particle Mesh Ewald (PME) method was used for long-range vdW interactions (83). The LINCS algorithm was applied to all bonds and angles of water molecules. The convergence of the simulations was monitored by plotting the toot-mean-square deviation of backbone atoms of the transmembrane helices of receptors and Gα subunits from the starting structure over time (Fig. S1a).

### Calculating the fingerprints of pairwise interactions between GPCR and Gα Protein

We utilized the Python script library “GetContacts” (https://www.github.com/getcontacts) to evaluate the nature of pairwise intermolecular residue contacts between GPCRGPCR and the G proteins. This script allows the identification of various interaction types, including salt bridges (cutoff <4.0 Å between anion and cation atoms), hydrogen bonds (cutoff <3.5 Å between hydrogen donor and acceptor atoms and an angle <70° between donor and acceptor), van der Waals interactions (a difference <2 Å between two atoms), pi-stack contacts (a distance <7.0 Å between the aromatic centers of residues and an angle <30° between the normal vectors emanating from each aromatic plane), and cation-pi contacts (a distance <6.0 Å between a cation atom and the centroid of an aromatic ring and an angle <60° between the normal vector from the aromatic plane to the cation atom). All types of contacts were considered in this study, and the contact analysis was performed on the compiled 5 μs trajectory, in which the water and ions were subsequently eliminated. The atom selection groups (‘--sele’ and ‘--sele2’) were matched with the corresponding amino acid residues for each protein domain, either GPCR or α-subunit of G protein. Each GPCR and G protein pdb number was converted into a corresponding generic residue number based on the BW numbering and the Common G protein numbering. Transmembrane helices ends were adapted based on the BW numbering for all receptors, except for β_2_AR: two present residues of ICL3 were given BW numbering of 5×77 and 5×78. Custom Python scripts were used to conduct one-hot encoding, generating a binary fingerprint for each simulation, where “1” represents the presence of a contact between two residues in a specific frame, and “0” indicates its absence.

### Bayesian network analysis of GPCR:G protein interactions

The binary fingerprints of the residue contact pairs were assessed to decipher their interdependent interactions using BNOmics, a tool developed for generalist BN analysis. Independent BNs were initially constructed for each GPCR:G protein complex. A search-and-score network model selection was executed using MU scoring function with 50 random restarts to ensure convergence (46). In the BN of contact fingerprints, the residue pairs serve as nodes, and the edge weight, which is a measure of variable dependency, is proportional to the ratio of the marginal maximum likelihood of the network with the edge to the one without, given the data. Notably, the novel MU scoring function/criterion is bound to a universal scale, thus allowing both intra– and inter-network comparisons (46). The network property of node strength, which is the total sum of edge weights related to this node, was used as an indicator of contact pairs’ connectivity. After arranging the residue pairs from the highest to the lowest node strength, the top 25 percent were compared across different G protein subtypes. The network visualization software Cytoscape 3.9.1 (84) was used to graphically represent these nodes and their interconnections.

### Robustness assessment of Bayesian network models

To rigorously assess the robustness of the BN models generated by BNOmics, we employed a two-pronged approach: randomization tests and sensitivity experiments. Randomization Tests: For each protein complex network model, we introduced 1000 small local random perturbations. These perturbations were designed to simulate small changes in the network topology by randomly adding and/or removing edges. The perturbed models were then scored using the same scoring functions employed by BNOmics (MU). These scores were compared against the score of the original, unperturbed model (Fig.S3a). Sensitivity Tests: We conducted a series of sensitivity experiments by randomly resampling the input MDs trajectory frames. The original dataset for each GPCR:G protein complex was randomly resampled 1000 times with replacement. Using the resampled data, we recalculated the scores for the new data, given the original network topologies. The rescored models were then evaluated to determine if their scores fell within the 0.99 confidence interval (Fig.S3b).

### Directed graph moralization

The directed graphs obtained in BNOmics were converted into moralized graphs using following steps: 1) for each node in the directed graph, its parent nodes were identified; 2) for each set of parent nodes, an undirected edge was added between every pair of parents that are not already connected; 3) all directed edges in the graph were converted to undirected edges. The algorithm ensures that the resulting moral graph is undirected and that all original dependencies are preserved. However, even though edges in the BNs are weighted, we have considered all edges as unweighted in the moralized graphs. The Python library NetworkX was employed for graph manipulation and analysis.

### Jensen-Shannon Divergence matrix of joint probabilities of the largest common subgraph

Using the moralization algorithm described prior, we obtained the largest common subnetwork (*G\**) between 5 complexes out of 6. Given *G\** and 5 datasets corresponding to the protein complexes, we calculated joint probability *J_k_* directly from the observation in each dataset *k*. We then used Jensen-Shannon Divergence (JSD) to compare the complexes. JSD was calculated between *J_i_* and *J_k_*, where *i* and *k* are two datasets, pairwise between all complexes and results were represented as heatmap (Fig. 4c).

## Supplemental Figure Legends

**Supplemental Figure 1.**
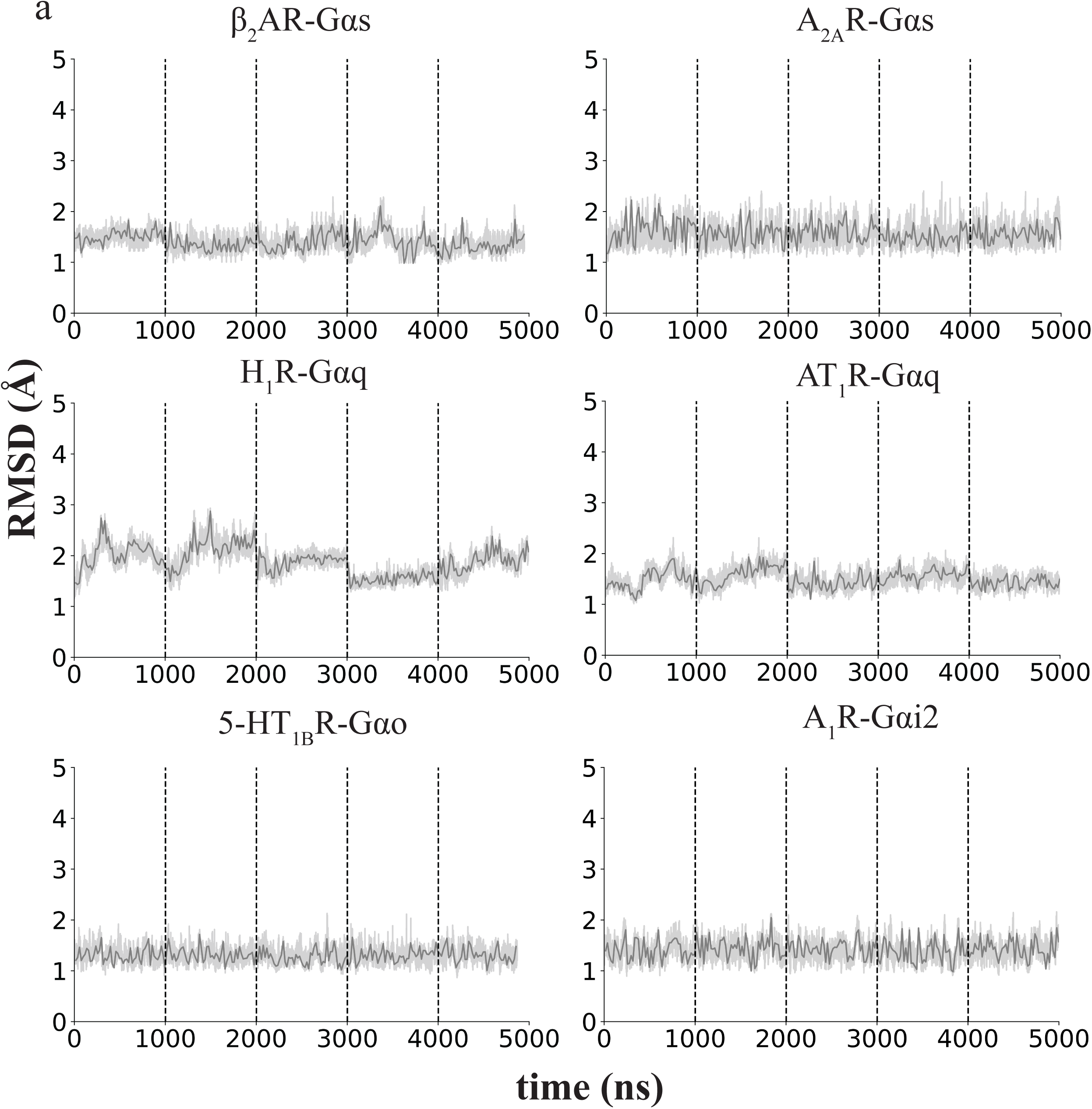
**a**. Root-mean-square deviation (RMSD) values for TM backbone atoms in the transmembrane helices.

**Supplemental Figure 2.**
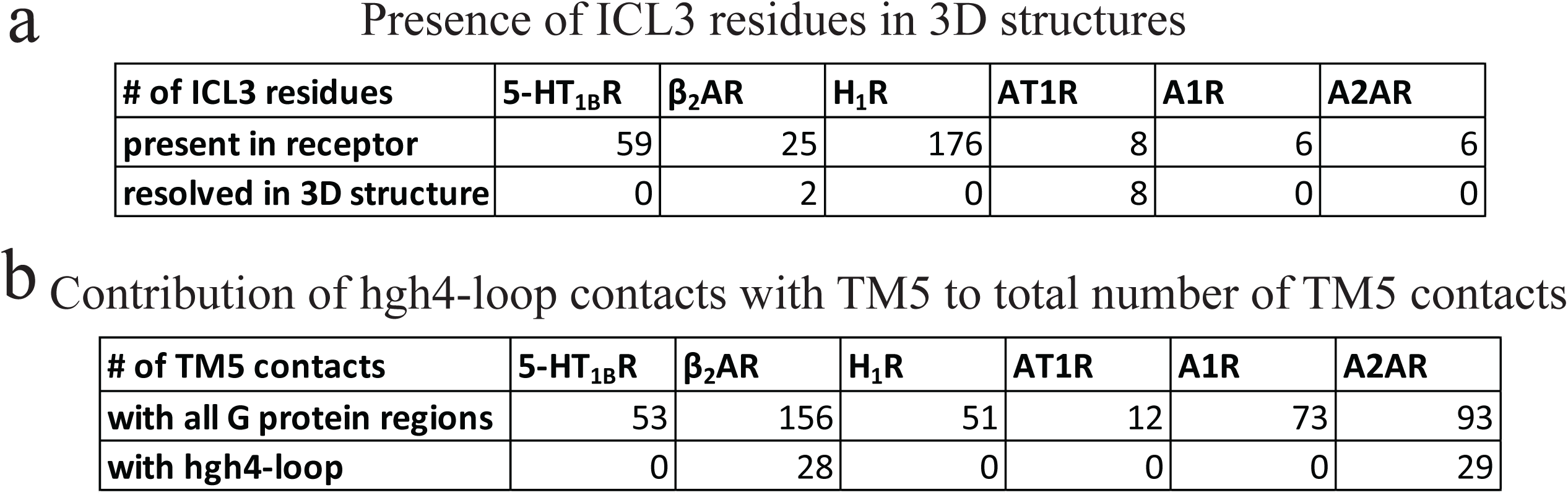
**a**. Table demonstrating the total number of ICL3 residues across GPCRs analyzed in this study and the number of resolved residues in the three-dimensional structures used for the MD simulations. **b.** Table demonstrating the total number of TM5 interactions with any G protein region and the number of contacts that TM5 makes only with hgh4-loop of G protein core region.

**Supplemental Figure 3.**
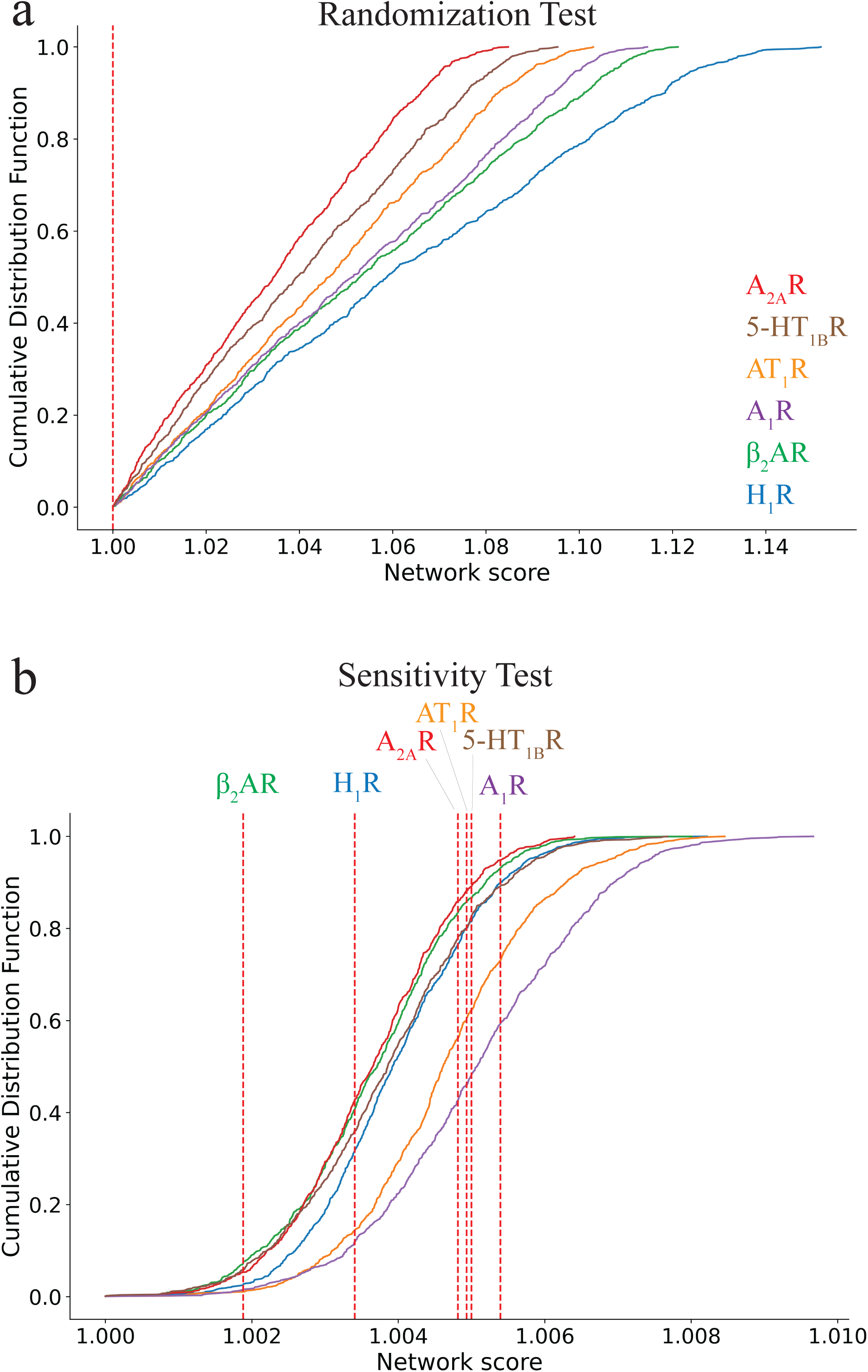
Evaluation of Bayesian Network Analysis Robustness. **a.** Cumulative Distribution Function of all network scores with various topology perturbations. Red dashed line indicates the placement of score of the original networks. **b.** Cumulative Distribution Function of all network scores with data resampling. Red dashed line indicates the placement of score of the original networks.

**Supplemental Figure 4.**
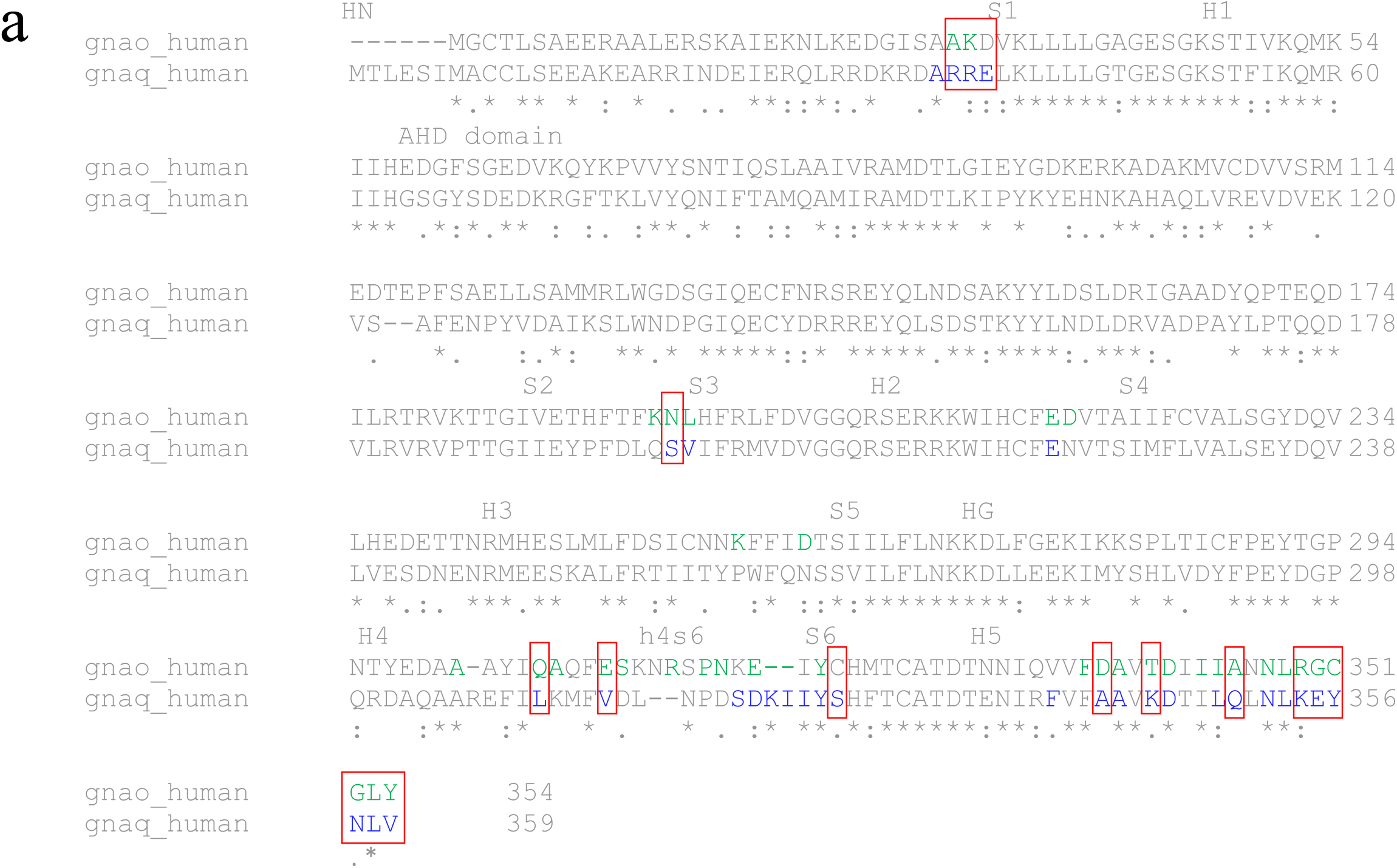
Alignment of Go and Gq protein sequences. **a.** BN-predicted cooperativity hotspots are marked in colored font according to the subtype: green – Go/Gi protein, blue – Gq protein. Selectivity hotspots are shown in red boxes.

## Acknowledgements

This work was funded by grants from the National Institutes of Health R01-GM117923 to N.V., R01-LM013876 to N.V., A.R., S.B., and S.Bh., and R01-LM013138 to A.R. Additional support is acknowledged from Dr. Susumu Ohno Chair in Theoretical Biology (held by A.R.), and Susumu Ohno Distinguished Investigator Fellowship (to G.G.).

## Author Contributions

N.V., A.R., S.B. conceptualized the project. E.M., N.M., and W.J.C.V performed the MD simulations. E.M. and S.Bh., did the analysis. G.G developed the code for BNOmics; N.V., A.R. and S.B. were involved in funding acquisition. E.M., N.V., A.R., N.M., S.B. and S.Bh. were involved in writing, reviewing, and editing of the manuscript.

## References

1. Armstrong, J. F., Faccenda, E., Harding, S. D., Pawson, A. J., Southan, C., Sharman, J. L., Campo, B., Cavanagh, D. R., Alexander, S. P. H., Davenport, A. P., Spedding, M., Davies, J. A., and Nc, I. (2020) The IUPHAR/BPS Guide to PHARMACOLOGY in 2020: extending immunopharmacology content and introducing the IUPHAR/MMV Guide to MALARIA PHARMACOLOGY. Nucleic Acids Res 48, D1006–D1021

2. Masuho, I., Ostrovskaya, O., Kramer, G. M., Jones, C. D., Xie, K., and Martemyanov, K. A. (2015) Distinct profiles of functional discrimination among G proteins determine the actions of G protein-coupled receptors. Sci Signal 8, ra123-ra123

3. Lukasheva, V., Devost, D., Le Gouill, C., Namkung, Y., Martin, R. D., Longpré, J.-M., Amraei, M., Shinjo, Y., Hogue, M., Lagacé, M., Breton, B., Aoki, J., Tanny, J. C., Laporte, S. A., Pineyro, G., Inoue, A., Bouvier, M., and Hébert, T. E. (2020) Signal profiling of the β(1)AR reveals coupling to novel signalling pathways and distinct phenotypic responses mediated by β(1)AR and β(2)AR. Sci Rep 10, 8779–8779

4. Okashah, N., Wan, Q., Ghosh, S., Sandhu, M., Inoue, A., Vaidehi, N., and Lambert, N. A. (2019) Variable G protein determinants of GPCR coupling selectivity. Proc Natl Acad Sci U S A 116, 12054–12059

5. Sandhu, M., Cho, A., Ma, N., Mukhaleva, E., Namkung, Y., Lee, S., Ghosh, S., Lee, J. H., Gloriam, D. E., Laporte, S. A., Babu, M. M., and Vaidehi, N. (2022) Dynamic spatiotemporal determinants modulate GPCR:G protein coupling selectivity and promiscuity. Nat Commun 13, 7428–7428

6. Sandhu, M., Touma, A. M., Dysthe, M., Sadler, F., Sivaramakrishnan, S., and Vaidehi, N. (2019) Conformational plasticity of the intracellular cavity of GPCR-G-protein complexes leads to G-protein promiscuity and selectivity. Proc Natl Acad Sci U S A 116, 11956–11965

7. Semack, A., Sandhu, M., Malik, R. U., Vaidehi, N., and Sivaramakrishnan, S. (2016) Structural Elements in the Gαs and Gαq C Termini That Mediate Selective G Protein-coupled Receptor (GPCR) Signaling. J Biol Chem 291, 17929–17940

8. Masuho, I., Ostrovskaya, O., Kramer, G. M., Jones, C. D., Xie, K., and Martemyanov, K. A. Distinct profiles of functional discrimination among G proteins determine the actions of G protein-coupled receptors.

9. Wenzel-Seifert, K., and Seifert, R. Molecular analysis of beta(2)-adrenoceptor coupling to G(s)-, G(i)-, and G(q)-proteins.

10. Stallaert, W., Dorn J f Fau – van der Westhuizen, E., van der Westhuizen E Fau – Audet, M., Audet M Fau – Bouvier, M., and Bouvier, M. Impedance responses reveal β₂-adrenergic receptor signaling pluridimensionality and allow classification of ligands with distinct signaling profiles.

11. Malik, R. U., Ritt M Fau – DeVree, B. T., DeVree Bt Fau – Neubig, R. R., Neubig Rr Fau – Sunahara, R. K., Sunahara Rk Fau – Sivaramakrishnan, S., and Sivaramakrishnan, S. Detection of G protein-selective G protein-coupled receptor (GPCR) conformations in live cells.

12. Semack, A., Malik, R. U., and Sivaramakrishnan, S. G Protein-selective GPCR Conformations Measured Using FRET Sensors in a Live Cell Suspension Fluorometer Assay. LID – 10.3791/54696 [doi] LID – 54696.

13. Mackenzie, A. E., Quon, T., Lin, L. C., Hauser, A. S., Jenkins, L., Inoue, A., Tobin, A. B., Gloriam, D. E., Hudson, B. D., and Milligan, G. Receptor selectivity between the G proteins Gα(12) and Gα(13) is defined by a single leucine-to-isoleucine variation.

14. Olsen, R. H. J., DiBerto, J. F., English, J. G., Glaudin, A. M., Krumm, B. E., Slocum, S. T., Che, T., Gavin, A. C., McCorvy, J. D., Roth, B. L., and Strachan, R. T. (2020) TRUPATH, an open-source biosensor platform for interrogating the GPCR transducerome. Nature Chemical Biology 16, 841–849

15. Avet, C., Mancini, A., Breton, B., Le Gouill, C., Hauser, A. S., Normand, C., Kobayashi, H., Gross, F., Hogue, M., Lukasheva, V., St-Onge, S., Carrier, M., Héroux, M., Morissette, S., Fauman, E. B., Fortin, J.-P., Schann, S., Leroy, X., Gloriam, D. E., and Bouvier, M. (2022) Effector membrane translocation biosensors reveal G protein and βarrestin coupling profiles of 100 therapeutically relevant GPCRs. Elife 11, e74101

16. Inoue, A., Raimondi, F., Kadji, F. M. N., Singh, G., Kishi, T., Uwamizu, A., Ono, Y., Shinjo, Y., Ishida, S., Arang, N., Kawakami, K., Gutkind, J. S., Aoki, J., and Russell, R. B. Illuminating G-Protein-Coupling Selectivity of GPCRs.

17. Glukhova, A., Draper-Joyce, C. J., Sunahara, R. K., Christopoulos, A., Wootten, D., and Sexton, P. M. Rules of Engagement: GPCRs and G Proteins.

18. Kim, H. A.-O. X., Xu, J., Maeda, S. A.-O., Duc, N. A.-O., Ahn, D. A.-O., Du, Y. A.-O., and Chung, K. A.-O. Structural mechanism underlying primary and secondary coupling between GPCRs and the Gi/o family.

19. Wess, J., Liu J Fau – Blin, N., Blin N Fau – Yun, J., Yun J Fau – Lerche, C., Lerche C Fau – Kostenis, E., and Kostenis, E. Structural basis of receptor/G protein coupling selectivity studied with muscarinic receptors as model systems.

20. Alegre, K. O., Paknejad, N., Su, M., Lou, J.-S., Huang, J., Jordan, K. D., Eng, E. T., Meyerson, J. R., Hite, R. K., and Huang, X.-Y. (2021) Structural basis and mechanism of activation of two different families of G proteins by the same GPCR. Nature Structural & Molecular Biology 28, 936–944

21. Harris, J. A., Faust, B., Gondin, A. B., Dämgen, M. A., Suomivuori, C.-M., Veldhuis, N. A., Cheng, Y., Dror, R. O., Thal, D. M., and Manglik, A. (2022) Selective G protein signaling driven by substance P– neurokinin receptor dynamics. Nature Chemical Biology 18, 109–115

22. Suno, R., Sugita, Y., Morimoto, K., Takazaki, H., Tsujimoto, H., Hirose, M., Suno-Ikeda, C., Nomura, N., Hino, T., Inoue, A., Iwasaki, K., Kato, T., Iwata, S., and Kobayashi, T. Structural insights into the G protein selectivity revealed by the human EP3-G(i) signaling complex.

23. Huang, S., Xu, P., Shen, D. D., Simon, I. A., Mao, C., Tan, Y., Zhang, H., Harpsøe, K., Li, H., Zhang, Y., You, C., Yu, X., Jiang, Y., Zhang, Y., Gloriam, D. E., and Xu, H. E. GPCRs steer G(i) and G(s) selectivity via TM5-TM6 switches as revealed by structures of serotonin receptors.

24. Liu, Q., Yang, D., Zhuang, Y., Croll, T. I., Cai, X., Dai, A., He, X., Duan, J., Yin, W., Ye, C., Zhou, F., Wu, B., Zhao, Q., Xu, H. E., Wang, M.-W., and Jiang, Y. (2021) Ligand recognition and G-protein coupling selectivity of cholecystokinin A receptor. Nature Chemical Biology 17, 1238–1244

25. Mobbs, J. I., Belousoff, M. J., Harikumar, K. G., Piper, S. J., Xu, X., Furness, S. G. B., Venugopal, H., Christopoulos, A., Danev, R., Wootten, D., Thal, D. M., Miller, L. J., and Sexton, P. M. (2021) Structures of the human cholecystokinin 1 (CCK1) receptor bound to Gs and Gq mimetic proteins provide insight into mechanisms of G protein selectivity. PLOS Biology 19, e3001295

26. del Sol, A., Fujihashi, H., Amoros, D., and Nussinov, R. (2006) Residues crucial for maintaining short paths in network communication mediate signaling in proteins. Molecular Systems Biology 2, 2006.0019

27. Hilser, V. J., Dowdy D Fau – Oas, T. G., Oas Tg Fau – Freire, E., and Freire, E. The structural distribution of cooperative interactions in proteins: analysis of the native state ensemble.

28. Prabhakar, P. A.-O., Ray, D. A.-O., and Andricioaei, I. A.-O. Predicting residue cooperativity during protein folding: A combined, molecular dynamics and unsupervised learning approach.

29. Moza, B., Buonpane, R. A., Zhu, P., Herfst, C. A., Rahman, A. K. M. N.-u., McCormick, J. K., Kranz, D. M., and Sundberg, E. J. (2006) Long-range cooperative binding effects in a T cell receptor variable domain. Proceedings of the National Academy of Sciences 103, 9867–9872

30. de Vink, P. A.-O., Andrei, S. A.-O., Higuchi, Y. A.-O., Ottmann, C. A.-O., Milroy, L. A.-O., and Brunsveld, L. A.-O. X. Cooperativity basis for small-molecule stabilization of protein-protein interactions.

31. Park, D., Izaguirre, J., Coffey, R., and Xu, H. (2023) Modeling the Effect of Cooperativity in Ternary Complex Formation and Targeted Protein Degradation Mediated by Heterobifunctional Degraders. ACS Bio & Med Chem Au 3, 74–86

32. Pearl, J. (1988) Chapter 1 – UNCERTAINTY IN AI SYSTEMS: AN OVERVIEW. in Probabilistic Reasoning in Intelligent Systems (Pearl, J. ed.), Morgan Kaufmann, San Francisco (CA). pp 1-28

33. Pearl, J. (2000) Causality: models, reasoning, and inference, Cambridge University Press

34. Bhattacharya, S., Salomon-Ferrer, R., Lee, S., and Vaidehi, N. (2016) Conserved Mechanism of Conformational Stability and Dynamics in G-Protein-Coupled Receptors. J Chem Theory Comput 12, 5575–5584

35. Bhattacharya, S., and Vaidehi, N. (2014) Differences in allosteric communication pipelines in the inactive and active states of a GPCR. Biophys J 107, 422–434

36. Krumm, B. E., Lee, S., Bhattacharya, S., Botos, I., White, C. F., Du, H., Vaidehi, N., and Grisshammer, R. (2016) Structure and dynamics of a constitutively active neurotensin receptor. Sci Rep 6, 38564–38564

37. Lee, S., Bhattacharya, S., Tate, C. G., Grisshammer, R., and Vaidehi, N. (2015) Structural dynamics and thermostabilization of neurotensin receptor 1. J Phys Chem B 119, 4917–4928

38. Lee, S., Devamani, T., Song, H. D., Sandhu, M., Larsen, A., Sommese, R., Jain, A., Vaidehi, N., and Sivaramakrishnan, S. (2017) Distinct structural mechanisms determine substrate affinity and kinase activity of protein kinase Cα. J Biol Chem 292, 16300–16309

39. Lee, S., Nivedha, A. K., Tate, C. G., and Vaidehi, N. (2019) Dynamic Role of the G Protein in Stabilizing the Active State of the Adenosine A(2A) Receptor. Structure 27, 703–712.e703

40. Ma, N., Lee, S., and Vaidehi, N. (2020) Activation Microswitches in Adenosine Receptor A(2A) Function as Rheostats in the Cell Membrane. Biochemistry 59, 4059–4071

41. Ma, N., Lippert, L. G., Devamani, T., Levy, B., Lee, S., Sandhu, M., Vaidehi, N., and Sivaramakrishnan, S. (2018) Bitopic Inhibition of ATP and Substrate Binding in Ser/Thr Kinases through a Conserved Allosteric Mechanism. Biochemistry 57, 6387–6390

42. Gogoshin, G., Boerwinkle, E., and Rodin, A. S. (2017) New Algorithm and Software (BNOmics) for Inferring and Visualizing Bayesian Networks from Heterogeneous Big Biological and Genetic Data. J Comput Biol 24, 340–356

43. Rodin, A. S., Gogoshin, G., Hilliard, S., Wang, L., Egelston, C., Rockne, R. C., Chao, J., and Lee, P. P. (2021) Dissecting Response to Cancer Immunotherapy by Applying Bayesian Network Analysis to Flow Cytometry Data. International Journal of Molecular Sciences 22, 2316

44. Wang, X., Branciamore, S., Gogoshin, G., Ding, S., and Rodin, A. S. (2020) New Analysis Framework Incorporating Mixed Mutual Information and Scalable Bayesian Networks for Multimodal High Dimensional Genomic and Epigenomic Cancer Data. Front Genet 11, 648–648

45. Gogoshin, G., Branciamore, S., and Rodin, A. S. (2021) Synthetic data generation with probabilistic Bayesian Networks. Mathematical Biosciences and Engineering 18, 8603–8621

46. Gogoshin, G., and Rodin, A. S. (2023) Minimum Uncertainty as Bayesian Network Model Selection Principle. in *Under review, BMC Bioinformatics*, Preprint at 10.20944/preprints202202.0254.v2.

47. Doshi, U., Holliday, M. J., Eisenmesser, E. Z., and Hamelberg, D. (2016) Dynamical network of residue-residue contacts reveals coupled allosteric effects in recognition, catalysis, and mutation. Proc Natl Acad Sci U S A 113, 4735–4740

48. Johnson, Q. R., Lindsay, R. J., Nellas, R. B., Fernandez, E. J., and Shen, T. (2015) Mapping allostery through computational glycine scanning and correlation analysis of residue-residue contacts. Biochemistry 54, 1534–1541

49. Johnson, Q. R., Lindsay, R. J., and Shen, T. (2018) CAMERRA: An analysis tool for the computation of conformational dynamics by evaluating residue-residue associations. Journal of Computational Chemistry 39, 1568–1578

50. Zhou, Q., Yang, D., Wu, M., Guo, Y., Guo, W., Zhong, L., Cai, X., Dai, A., Jang, W., Shakhnovich, E. I., Liu, Z.-J., Stevens, R. C., Lambert, N. A., Babu, M. M., Wang, M.-W., and Zhao, S. (2019) Common activation mechanism of class A GPCRs. Elife 8, e50279

51. Kleino, I., Frolovaitė, P., Suomi, T., and Elo, L. L. (2022) Computational solutions for spatial transcriptomics. Comput Struct Biotechnol J 20, 4870–4884

52. Razavian, N. S., Kamisetty, H., and Langmead, C. J. (2012) Learning generative models of molecular dynamics. BMC Genomics 13, S5

53. Grzegorczyk, M. (2010) An Introduction to Gaussian Bayesian Networks. in Methods in Molecular Biology, Humana Press

54. Chen, L., Gong, W., Han, Z., Zhou, W., Yang, S., and Li, C. (2022) Key Residues in δ Opioid Receptor Allostery Explored by the Elastic Network Model and the Complex Network Model Combined with the Perturbation Method. Journal of Chemical Information and Modeling 62, 6727–6738

55. Lim Chong, W., Vao-soongnern, V., Nimmanpipug, P., Tayapiwatana, C., Lin, J.-H., Lin, Y.-L., Yee Chee, H., Md Zain, S., Abd Rahman, N., and Sanghiran Lee, V. (2022) Molecular dynamics simulations and Gaussian network model for designing antibody mimicking protein towards dengue envelope protein. Journal of Molecular Liquids 346, 118086

56. Rasmussen, S. G., DeVree Bt Fau – Zou, Y., Zou Y Fau – Kruse, A. C., Kruse Ac Fau – Chung, K. Y., Chung Ky Fau – Kobilka, T. S., Kobilka Ts Fau – Thian, F. S., Thian Fs Fau – Chae, P. S., Chae Ps Fau – Pardon, E., Pardon E Fau – Calinski, D., Calinski D Fau – Mathiesen, J. M., Mathiesen Jm Fau – Shah, S. T. A., Shah St Fau – Lyons, J. A., Lyons Ja Fau – Caffrey, M., Caffrey M Fau – Gellman, S. H., Gellman Sh Fau – Steyaert, J., Steyaert J Fau – Skiniotis, G., Skiniotis G Fau – Weis, W. I., Weis Wi Fau – Sunahara, R. K., Sunahara Rk Fau – Kobilka, B. K., and Kobilka, B. K. Crystal structure of the β2 adrenergic receptor-Gs protein complex.

57. Draper-Joyce, C. J., Khoshouei, M., Thal, D. M., Liang, Y. L., Nguyen, A. T. N., Furness, S. G. B., Venugopal, H., Baltos, J. A., Plitzko, J. M., Danev, R., Baumeister, W., May, L. T., Wootten, D., Sexton, P. M., Glukhova, A., and Christopoulos, A. Structure of the adenosine-bound human adenosine A(1) receptor-G(i) complex.

58. Xia, R., Wang, N., Xu, Z., Lu, Y., Song, J., Zhang, A., Guo, C., and He, Y. A.-O. X. Cryo-EM structure of the human histamine H(1) receptor/G(q) complex.

59. Zhang, D. A.-O., Liu, Y. A.-O., Zaidi, S. A., Xu, L., Zhan, Y. A.-O., Chen, A., Guo, J. A.-O. X., Huang, X. P., Roth, B. A.-O., Katritch, V. A.-O., Cherezov, V. A.-O., and Zhang, H. A.-O. Structural insights into angiotensin receptor signaling modulation by balanced and biased agonists.

60. García-Nafría, J., Lee, Y., Bai, X., Carpenter, B. A.-O., and Tate, C. A.-O. Cryo-EM structure of the adenosine A(2A) receptor coupled to an engineered heterotrimeric G protein. LID – 10.7554/eLife.35946 [doi] LID – e35946.

61. Selçuk, B. A.-O., Erol, I. A.-O., Durdağı, S. A.-O., and Adebali, O. A.-O. Evolutionary association of receptor-wide amino acids with G protein-coupling selectivity in aminergic GPCRs. LID – 10.26508/lsa.202201439 [doi] LID – e202201439.

62. Ballesteros, J. A., and Weinstein, H. (1995) [19] Integrated methods for the construction of three-dimensional models and computational probing of structure-function relations in G protein-coupled receptors. in Methods in Neurosciences (Sealfon, S. C. ed.), Academic Press. pp 366–428

63. Flock, T., Ravarani, C. N. J., Sun, D., Venkatakrishnan, A. J., Kayikci, M., Tate, C. G., Veprintsev, D. B., and Babu, M. M. Universal allosteric mechanism for Gα activation by GPCRs.

64. Sadler, F., Ma, N., Ritt, M., Sharma, Y., Vaidehi, N., and Sivaramakrishnan, S. (2023) Autoregulation of GPCR signalling through the third intracellular loop. Nature 615, 734–741

65. Jelinek, V. A.-O., Mösslein, N. A.-O., and Bünemann, M. A.-O. Structures in G proteins important for subtype selective receptor binding and subsequent activation.

66. Flock, T., Hauser, A. S., Lund, N., Gloriam, D. E., Balaji, S., and Babu, M. M. (2017) Selectivity determinants of GPCR-G-protein binding. Nature 545, 317–322

67. Han, J., Zhang, J., Nazarova, A. A.-O., Bernhard, S. M., Krumm, B. A.-O., Zhao, L., Lam, J. H., Rangari, V. A., Majumdar, S. A.-O., Nichols, D. E., Katritch, V. A.-O., Yuan, P. A.-O. X., Fay, J. A.-O., and Che, T. A.-O. Ligand and G-protein selectivity in the κ-opioid receptor.

68. Kling, R. C., Lanig, H., Clark, T., and Gmeiner, P. (2013) Active-State Models of Ternary GPCR Complexes: Determinants of Selective Receptor-G-Protein Coupling. PLOS ONE 8, e67244

69. Zhao, L. A.-O., Lin, J. A.-O., Ji, S. Y., Zhou, X. E., Mao, C. A.-O., Shen, D. A.-O., He, X., Xiao, P. A.-O., Sun, J. A.-O., Melcher, K., Zhang, Y. A.-O., Yu, X. A.-O., and Xu, H. A.-O. Structure insights into selective coupling of G protein subtypes by a class B G protein-coupled receptor.

70. McGlohon, M. (2007) Methods and Uses of Graph Demoralization.

71. Franziska, M. H., Maria, M.-S., Manbir, S., Brian, K. K., Michel, B., and Babu, M. M. (2021) Dissecting the allosteric networks governing agonist efficacy and potency in G protein-coupled receptors. bioRxiv, 2021.2009.2014.460253

72. Neumann, S., Krause G Fau – Claus, M., Claus M Fau – Paschke, R., and Paschke, R. Structural determinants for g protein activation and selectivity in the second intracellular loop of the thyrotropin receptor.

72. Venkatakrishnan, A. J., Fonseca, R., Ma, A. K., Hollingsworth, S. A., Chemparathy, A., Hilger, D., Kooistra, A. J., Ahmari, R., Babu, M. M., Kobilka, B. K., and Dror, R. O. (2019) Uncovering patterns of atomic interactions in static and dynamic structures of proteins. Cold Spring Harbor Laboratory

74. Tsai, C.-J., Marino, J., Adaixo, R., Pamula, F., Muehle, J., Maeda, S., Flock, T., Taylor, N. M., Mohammed, I., Matile, H., Dawson, R. J., Deupi, X., Stahlberg, H., and Schertler, G. (2019) Cryo-EM structure of the rhodopsin-Gαi-βγ complex reveals binding of the rhodopsin C-terminal tail to the gβ subunit. Elife 8, e46041

75. Vanommeslaeghe, K., Hatcher E Fau – Acharya, C., Acharya C Fau – Kundu, S., Kundu S Fau – Zhong, S., Zhong S Fau – Shim, J., Shim J Fau – Darian, E., Darian E Fau – Guvench, O., Guvench O Fau – Lopes, P., Lopes P Fau – Vorobyov, I., Vorobyov I Fau – Mackerell, A. D., Jr., and Mackerell, A. D., Jr. CHARMM general force field: A force field for drug-like molecules compatible with the CHARMM all-atom additive biological force fields.

76. Lomize, M. A., Pogozheva Id Fau – Joo, H., Joo H Fau – Mosberg, H. I., Mosberg Hi Fau – Lomize, A. L., and Lomize, A. L. OPM database and PPM web server: resources for positioning of proteins in membranes.

77. Jo, S., Lim Jb Fau – Klauda, J. B., Klauda Jb Fau – Im, W., and Im, W. CHARMM-GUI Membrane Builder for mixed bilayers and its application to yeast membranes.

78. Jo S Fau – Kim, T., Kim T Fau – Iyer, V. G., Iyer Vg Fau – Im, W., and Im, W. CHARMM-GUI: a web-based graphical user interface for CHARMM.

78. Huang, J., Rauscher, S., Nawrocki, G., Ran, T., Feig, M., de Groot, B. L., Grubmüller, H., and MacKerell, A. D. (2017) CHARMM36m: an improved force field for folded and intrinsically disordered proteins. Nature Methods 14, 71–73

80. Abraham, M. J., Murtola, T., Schulz, R., Páll, S., Smith, J. C., Hess, B., and Lindahl, E. (2015) GROMACS: High performance molecular simulations through multi-level parallelism from laptops to supercomputers. SoftwareX 1-2, 19–25

81. M.J. Abraham, D. v. d. S. E. Lindahl, B. Hess, and the GROMACS development team. GROMACS User Manual version 2019.

82. Parrinello, M., and Rahman, A. (1981) Polymorphic transitions in single crystals: A new molecular dynamics method. Journal of Applied Physics 52, 7182–7190

83. Darden, T., York, D., and Pedersen, L. (1993) Particle mesh Ewald: An N⋅log(N) method for Ewald sums in large systems. The Journal of Chemical Physics 98, 10089–10092

84. Franz, M., Lopes, C. T., Huck, G., Dong, Y., Sumer, O., and Bader, G. D. (2016) Cytoscape.js: a graph theory library for visualisation and analysis. Bioinformatics 32, 309–311

